# Experimental corticosterone manipulation increases mature feather corticosterone content: implications for inferring avian stress history from feather analyses

**DOI:** 10.1101/2021.01.11.425815

**Authors:** Yaara Aharon-Rotman, William A. Buttemer, Lee Koren, Katherine Wynne-Edwards

## Abstract

Feathers incorporate plasma corticosterone (CORT) during their development and, because feathers lack blood supply at maturity, the CORT content of feathers (CORTf) is presumed to represent an integrated average of plasma CORT levels during feather growth. We tested this assumption by quantifying CORTf in feathers of house sparrows (Passer domesticus) that were plucked before and after being implanted with either a corticosterone-filled, metyrapone-filled, or empty (sham) silastic capsule. Two of the seven flight feathers collected from each bird were fully grown when birds received implants. We found that CORTf of all seven feathers corresponded with treatment type, and with plasma CORT levels of non-moulting reference sparrows also given these implants. We also found that CORTf of the two mature feathers of each bird were 4 to 10-fold higher than values measured in new feathers. Given the avascular nature of mature feathers, and the fact that we did not wash the feathers prior to analysis, the most plausible explanation for our results is that CORT was externally deposited on feathers after implant. This outcome emphasises the need for follow-up studies to identify the external sources of CORT that may affect the CORTf of feathers. We hope this study will stimulate discussions and further much needed studies on the mechanism of CORT deposition in feathers and open exciting opportunities for application of such methods in ecological research, such as measuring multiple time scales in such a non-invasive manner.

## INTRODUCTION

Corticosterone (CORT) is the main glucocorticoid hormone in birds and studies have shown that circulating CORT is incorporated into developing feathers via diffusion from the blood during cell differentiation (Jenni-Eiermann *et al.* 2015). Feather CORT content (CORT_f_) thus varies with plasma CORT levels during feather replacement (Lattin *et al.* 2011; Fairhurst *et al.* 2013) and measurements of CORT_f_ are presumed to represent an integrated average of plasma CORT levels experienced by birds over the feather’s maturation period (Bortolotti *et al.* 2008; Romero and Fairhurst 2016). Consequently, CORT_f_ has gained prominence as a measure of hormone levels experienced by birds over the moulting period (Romero and Fairhurst 2016). However, studies are still required to answer questions regarding the use of feathers to reflect long-term plasma glucocorticoid levels (Sheriff *et al.* 2011). If the assumption that CORT_f_ represents circulating levels of CORT within only the period of feather development is, in fact, correct, experimental manipulation of CORT outside the moult period should have no effect on CORT_f_ of mature feathers.

The seminal study of CORT_f_ in red-legged partridges *Alectoris rufa* by Bortolotti *et al.* (2008) recognised that the lipophilic property of CORT could result in it being transferred to feather surfaces via preen oil, but discounted this possibility after finding CORT_f_ to be unaffected by cleansing feathers with a detergent solution. This conclusion was further supported by a study that could not detect CORT in the preen oil of starlings (*Sturnus vulgaris*; Lattin *et al.* 2011) and another which detected CORT in only one out of 23 uropygial oil samples from captive rhinoceros auklets (Cerorhinca monocerata; Will *et al.* 2019). These outcomes have resulted in the common practice to not consider the possibility for surface deposition of CORT onto mature feathers. However, experiments by Jenni-Eiermann *et al.* (2015) showed that washing feathers with a dilute detergent solution significantly reduced CORT_f_ of replacement feathers of captive feral pigeons (*Columba livia domestica*) receiving CORT-enriched pellets, but had no effect on replacement feathers CORT_f_ of control birds. This raises a critical question: are surface additions of CORT onto feathers of general concern for studies examining CORT_f_ in free-living birds, or are they an artefact of experiments producing super-physiological elevations of CORT during feather growth?

The aim of our study is to test the assumption that CORT is incorporated in feathers only during the development period. To this end, we quantified the effect of experimental manipulation of CORT on CORT_f_ of both fully grown (mature) and developing feathers in captive house sparrows (*Passer domesticus*). If indeed CORT is incorporated into feathers only during the development period, we expected the CORT_f_ of mature feathers to be unaffected by hormonal manipulation. However, if surface deposition does occur, CORT_f_ of both developing and mature feathers will vary with hormonal manipulation.

## MATERIALS AND METHODS

The description of the experimental methods and statistical analyses are detailed in the Supplementary Information. Briefly, we plucked 12 feathers of house sparrows after 1 week of adjustment to captivity. Only the six feathers plucked from the right wing were used in this study. Once feathers started to regrow (12 days after plucking), we manipulated corticosterone levels by implanting each bird with either (1) CORT-filled implants to achieve hyper-physiological CORT levels (N=7), (2) metyrapone-filled implants to attain hypo-physiological levels (N=9), and (3) empty implants (sham group) to sustain physiological levels (N=7). We validated the efficacy of the implants by measuring plasma CORT in a reference group of non-moulting sparrows given the same implants. Plasma CORT of the CORT-treated reference birds was persistently at hyper-physiological levels, initially low and then at hyper-physiological levels in the metyrapone-treated group, and remained at physiological levels in sham-treated birds (details in Supplementary Information). Upon completion of feather regrowth, ten replacement feathers were plucked, five to be used in this study (Fig. 1B) and 5 analysed and reported elsewhere (Aharon-Rotman *et al.* 2017).

**Figure 1:**
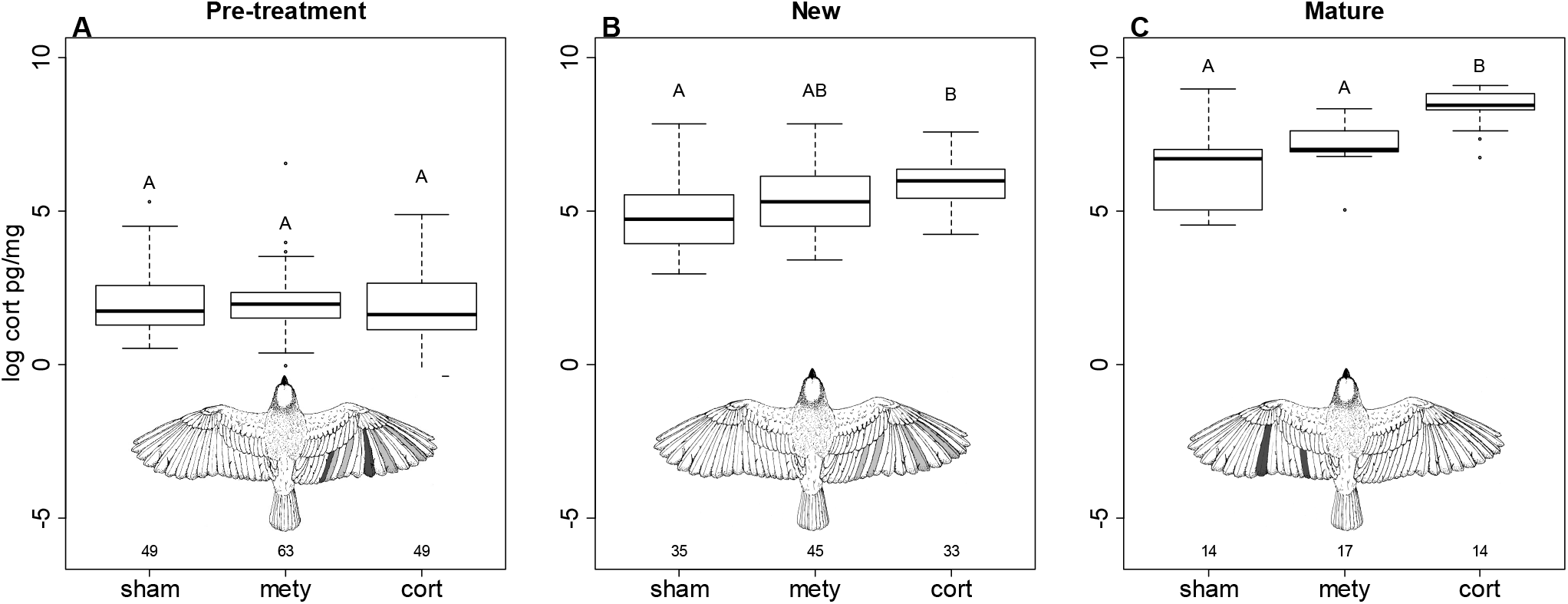
Boxplot of log corticosterone (cort) values in pg/mg feather in old feathers pre-treatment (“Pre-treatment”), re-grown feathers post treatment (“New”) and old feathers post-treatment (“Mature”), in relation to type of implant: sham, metyrapone (“mety”) and corticosterone (“cort”). Bird sketch in each panel depict the feathers used for this analysis. Note feathers in dark grey in the “pre treatment” (S5 and P2) were used as a reference to CORT_f_ values in “Mature” feathers (also in dark grey). Numbers below each bar depict the number of feathers in the analysis. Total number of individuals=23. Solid line depicts median values. Different letters above each box and whisker depict significant differences between treatments.

In addition, feathers P2 (primary) and S5 (secondary) from the left wing were plucked only after hormonal manipulation to determine if CORT_f_ of feathers that were mature throughout the experiment were affected by the treatment (Fig. 1C). We inferred their pre-treatment CORT content from CORT_f_ of complementary feathers P2 and S5 from the right wing that were plucked prior to the experiment (Fig. 1A, dark grey). Because birds lose and replace flight feathers in bilateral symmetry during moult, corresponding flight feathers from each wing share temporal development and should, accordingly, experience the same levels of CORT during their development. Comparison of CORT_f_ in corresponding flight feathers of several species confirms this assumption (e.g. Lattin *et al.* 2011). Thus, this study is based on a dataset of 7 “pre-treatment” feathers, 5 new replacement feathers (“new”) and 2 old feathers (“mature”) which were fully grown during hormonal manipulation and were plucked after the experiment (Fig. 1).

## RESULTS

We found a significant effect of treatment type (F=3.46_2,19_, *p*=*0.05*) on CORT_f_ values in the new, replacement feathers, which grew during hormonal manipulation. We also found a significant effect of treatment type (F=10.64_2,20_, *p*<*0.001*) on CORT_f_ values in the old feathers, which were mature during hormonal manipulation. Post-hoc tests revealed significant differences between CORT_f_ values of the sham and CORT-treated birds (t=−2.63, *p*=*0.04*) in the new feather group. In the mature feather group, differences were significant between both sham and metyrapone-treated birds compared with CORT-treated sparrows (t=−4.6, *p*<*0.001* and t=−2.72, *p*=*0.03*), respectively (Fig. 1 and Supplementary Table A1). A summary of the raw values is presented in Supplementary Table A2.

## DISCUSSION

We have demonstrated for the first time that feather corticosterone content of mature feathers can be affected by a bird’s circulating CORT outside the moult period. If, as is generally assumed, CORT were only deposited during feather synthesis (Bortolotti *et al.* 2008), there should be no difference between fully grown flight feathers plucked before CORT manipulation versus corresponding flight feathers which were completely mature during CORT manipulation. Thus, the parallel increases in CORT_f_ of mature and replacement feathers following experimental treatment was unexpected, but even more surprising was that mature feathers attained significantly higher CORT_f_ levels than replacement feathers.

The addition of CORT to mature feathers via diffusion from plasma is not a possibility, as anatomical studies show that the growing feather undergoes a retrograde resorption as it matures, resulting in an avascular, non-living tissue at moult completion (Lillie 1940). Given this, and the fact that we did not wash feathers prior to analysis, the most plausible explanation for the significant CORT_f_ increases we found in mature feathers was a result of external deposition. However, the mechanisms for this transfer are yet to be identified. Although birds in our study shared an outdoor flight cage after receiving implants, thus raising the potential for faecal and/or feather sheath fragments being deposited as surface contamination, we consider that surface deposition was self-administered. This is based on our finding that the CORT_f_ in mature feathers always aligned among birds within each treatment group and their mean and median values corresponded to the expected rank order based on plasma CORT levels of reference sparrows receiving the same implants (see Table A2 and Fig. A1 in Supplementary Information)

One of the routine maintenance activities of birds involves preening their feathers through mandibulation. Preening is typically preceded by wiping the bill on the preen gland. Because steroids are highly lipophilic, preen oil is a potential source of extravascular CORT. Although previous studies did not find CORT in preen oil (Bortolotti *et al.* 2008; Lattin *et al.* 2011; Will *et al.* 2019), there may be an opportunity for preen oils to acquire cutaneous steroids during preening, because epidermal tissues surrounding the calamus are highly vascularised during feather growth. Such a possibility, however, requires experimental validation.

Irrespective of lacking evidence for this source of CORT, external steroid transfer during preening may explain the higher values of CORT_f_ we found in mature feathers in comparison to new replacement feathers. Presuming that grooming activities persisted throughout the post-implant period, mature feathers would present a larger surface area for a longer period than the emerging feathers. This creates a greater potential for fully grown feathers to accumulate surface deposits of CORT over the 3-week period following CORT manipulation. We did not test for CORT in preen oil, and given the role of preen oil as waterproofing agent this hypothesis invites further speculation.

The application of our findings to measurements of CORT_f_ in free-living birds remains uncertain. Our experimental procedure produced supra-physiological levels of CORT experienced by the CORT-implanted birds that may cause exceptional CORT excretion in preen oil or saliva, which do not resemble natural response. However, we have suggested a potential mechanism for circulating CORT surface deposition in mature feathers, which needs further investigation. Importantly, although two of our hormone treatments produced hyperphysiological CORT levels in reference birds, plasma values for birds receiving empty implants are a consequence of endogenous CORT secretion and could occur under free-living conditions. Indeed other studies have found house sparrows to be very sensitive to caging and to show elevated capture-stress responses even following 9-weeks in captivity (Love *et al.* (2017).

What is critically needed is to further investigate the routes of entry of CORT into feathers and to identify possible external sources of CORT deposition that could affect the CORT_f_ of mature feathers. We hope this letter will stimulate discussion and provoke establishment of effective procedures, both in terms of solvent and duration, for removing surface residues without extracting CORT from within the feather. Such investigations and validation of methodology will raise confidence when inferring stress history using these relatively non-invasive methods in free-living birds and open exciting opportunities for application of such methods in ecological research, such as measuring multiple time scales.

## ACKNOWLEDGEMENTS

The research was facilitated by infrastructure and operating funds from the Canadian Foundation for Innovation, Alberta Innovation and Advanced Education and the Faculty of Veterinary Medicine at the University of Calgary to KWE. We thank Rod Collins for advice on analgesia and anaesthetic procedures and Gerhard Körtner for the bird sketch in Figure 1. All experiments were approved by the Deakin University Animal Ethics Committee and were conducted in conformity with the NHMRC Australian Code of Practice for the Care and Use of Animals for Scientific Purposes.

