## Supplementary Information for "Experimental corticosterone manipulation increases mature feather corticosterone content: implications for inferring avian stress history from feather analyses"

### A. Materials and Methods

EXPERIMENTAL ANIMALS

We captured 23 House sparrows *Passer domesticus* (11 females, 12 males) at Werribee, (Victoria, Australia 37.9°S, 144.65°E) in autumn 2013. Details of captive conditions, feather plucking, and hormone manipulation methods are also detailed in Aharon-Rotman et al. (2017). Briefly, after at least 1 week of adjustment to captivity, birds were anaesthetised before plucking 12 feathers from their wings and tail (five primaries (P), five secondaries (S) and two rectrices (R)). Once feathers started to regrow (12 days after plucking), we manipulated corticosterone levels by implanting each bird with either a single corticosterone-filled (Sigma-Aldrich, Sydney, New South Wales, Australia; #27840, 7 birds), metyrapone-filled (Sigma-Aldrich; #M2696, 9 birds), or empty silastic capsule ("sham", 7 birds). A day after receiving implants, birds were transferred from their holding cages to outdoor flight cages (ca. 2 x 3 x 2.5 metres) until replacement feathers were fully regrown. Our experience in maintaining house sparrows found that feathers are least damaged when birds are housed in outdoor aviaries compared to being held in small indoor cages. Food and water were refreshed daily, but birds were otherwise undisturbed to minimise provoking stress responses. Upon completion of feather regrowth, ten replacement feathers were plucked, five to be used in this study and 5 analysed and reported elsewhere (Aharon-Rotman et al. 2017). In addition, feathers P2 and S5 from the left wing were plucked to determine the effect of treatment on the $CORT_f$ of feathers that were mature during the treatment (Fig. 1C). This resulted in our current study having a dataset of 7 old feathers ("pre-treatment"), 5 new replacement feathers ("new") and 2 old feathers, which were fully grown (mature) during hormonal manipulation and were plucked after the experiment (Fig. 1).

VALIDATION OF IMPLANTS- PLASMA CORT IN REFERENCE GROUP

To minimise disturbance and consequent stress-reactions of aviary-housed birds, we measured plasma CORT in a reference group of 15 non-moulting sparrows given the same implants to validate the efficacy of the implants. The details of plasma CORT assay are detailed in Aharon-Rotman et al. 2017. Briefly, the sparrows were held for 2 weeks in captivity (details of holding room are as described for sparrows used for this study prior to plucking) prior to implantation of one of the three implant types (sham, metyrapone or CORT). Treatments were randomly assigned to the birds. When sampled, two birds were removed from their shared cage and had a 120 µl blood sample collected within 3 minutes of first being disturbed. The CORT and metyrapone-treated birds were sampled at 2, 5, 10 and 15 days after receiving implants, whereas the control birds were sampled 5 and 10 days after implantation.

Plasma CORT measurements of the reference group showed persistent hyper-physiological levels in CORT-treated birds (average over the experiment period of 180± 36.6ng $ml^{-1}$), initially low and then hyper-physiological levels in the metyrapone-treated group (average 91.4±25.45 ng $ml^{-1}$), and sustained physiological levels in sham-treated birds (average 25.31±3.7 ng $ml^{-1}$; Fig. A1). The increased levels of CORT in the metyrapone-treated birds after the first 10 days of low CORT values may have been caused by the rapid depletion of the metyrapone from its implant. It is also possible that the rate of metyrapone diffusing from capsules was inadequate to inhibit CORT synthesis, but was sufficient to interact with other components of the HPA axis, such that there was a rebound in CORT synthesis once the metyrapone release had ceased.

### LC-MS/MS QUANTITATION

We used liquid chromatography coupled to tandem mass spectrometry to eliminate potential antibody cross-reactivity and ensure that only corticosterone is quantified. Details of the MS method are provided in section B below. Briefly, each feather had the calamus removed, was weighed, and then extracted in a 13 × 100 mm borosilicate tube in 5 mL of methanol pre-chilled to 4˚C (LC/MS grade, Fisher Scientific). Extractions occurred at 4˚C for 20 h. The calibration line used the area ratio between the sample and the matched deuterated IS, fit with 1/x weighting. All r2 were > 0.995. Low and high QC yielded 90% and 93% precision over 5 runs, with a coefficient of variation of 10% and 7%, respectively.

### STATISTICAL ANALYSIS

Before statistical analyses, concentrations in ng/mL were converted to pg/mg based on calamus-free feather weight before extraction, rather than length, following Will et al. (2019). We analysed the effect of treatment on $CORT_f$ in both regrown feathers ("new") and feathers which were fully grown at the time of receiving implants and were collected at the end of the experiment ("mature"). For this purpose, we ran a general linear mixed effect model with individual as random effect. At first, $CORT_f$ (pg/mg feather) was set as the dependent variable in relation to treatment (CORT, sham or metyrapone), $CORT_f$ of feathers plucked prior to implants ("pre"; to use as reference $CORT_f$ values), sex and feather type ("new" or "mature"). Reference $CORT_f$ (of the corresponding feather plucked prior the experiment) was entered as a covariate as individuals with high pre-treatment $CORT_f$ have been shown to display higher $CORT_f$ post treatment (Aharon-Rotman et al. 2017). Since CORT levels may differ between sexes (but see Koren et al. (2012)), we included sex in the model. The initial model contained the main effects and all possible interaction terms. To evaluate differences between treatment groups, we used estimated marginal means post hoc test for multiple comparisons with Tukey method for adjusted p-value. Because there was a significant interaction between feather type and treatment, suggesting a different slope for the different feather types ($F_{2,132}$= 3.51, $p$<0.05), we separated the models by feather type. We therefore report the values for two separate models, with $CORT_f$ of new and old feathers as the dependent variable, respectively. The explanatory variables remained the same: treatment (CORT, sham or metyrapone), $CORT_f$ of feathers plucked prior to implants ("pre") and sex.

Secondly, we analysed the relation between $CORT_f$ of new, replacement feathers and those of mature feathers that were plucked after the experiment. We again used a general linear mixed effect model with individual as a random effect. $CORT_f$ (pg/mg feather) in the replacement feathers ("new") was set as the dependent variable in relation to $CORT_f$ in the mature feathers and treatment (CORT, sham or metyrapone).

Variables were excluded from models using ANOVA Type III, based on a threshold significance level set to $p$=0.05. We confirmed the use of random effect in the models by comparing the AIC of the best model with and without random effect using the REML method (Zuur et al. 2009). R-function *lme* in R- package "nlme" was used to perform mixed effect models with the function *visreg* in the "visreg" package for visualization, and R-function *emmeans* in the R package "emmeans" was used to compute the post-hoc test.

All statistical analyses were conducted using R version 3.3.0 (R Development Core Team, 2019).

All sparrows retained their implants throughout the experimental period, with no evidence of broken skin at the site of the wound when implants were removed. All metyrapone implants were completely depleted when removed, whereas all CORT implants had considerable residual content.

**Figure A1:**

Plasma CORT levels (ng ml$^{-1}$) in the implant validation study for birds treated with corticosterone (CORT), metyrapone or empty (sham) implants. Plasma CORT was measured 4 times during the 15-day experiment in CORT and metyrapone-treated birds and twice in sham-treated individuals.

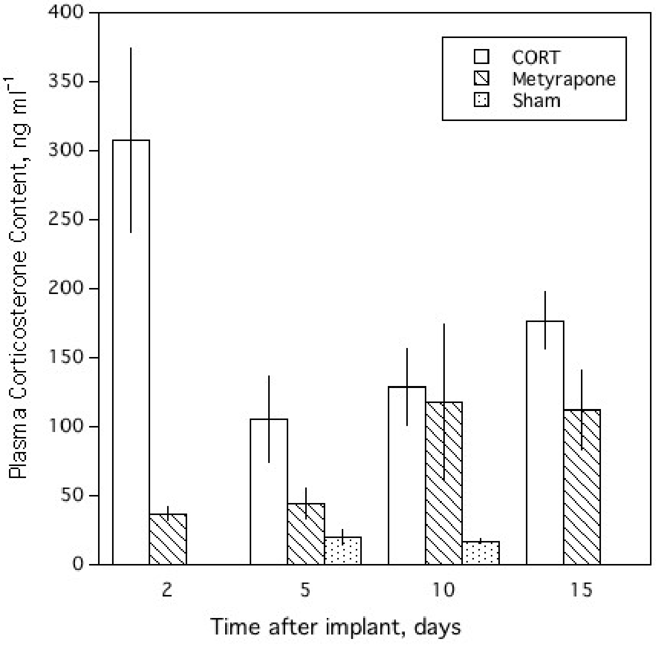

**Table A1:** Results of post-hoc estimated marginal means test for multiple comparisons with Tukey method for adjusted p-value. Post-hoc test was performed following a significant effect of treatment on corticosterone (cort) levels in feathers in both New and Mature feathers, using ANOVA type-3 test, to evaluate the differences between all treatment groups. Mety= metyrapone

| Contrast | Feather | Estimate | SE | df | t-ratio | p-value |
|---|---|---|---|---|---|---|
| sham-mety | New | -0.459 | 0.322 | 19 | -1.425 | 0.3489 |
| sham-cort | New | -0.911 | 0.346 | 19 | -2.631 | 0.0416 |
| mety-cort | New | -0.451 | 0.326 | 19 | -1.384 | 0.369 |
| sham-mety | Mature | -0.89 | 0.419 | 20 | -2.128 | 0.109 |
| sham-cort | Mature | -2.03 | 0.441 | 20 | -4.599 | 0.000 |
| mety-cort | Mature | -1.14 | 0.419 | 20 | -2.722 | 0.033 |

**Table A2:** Summary table (mean corticosterone (cort) values in pg/mg feather, median and standard deviation (SD)) for the two feather groups as measure before (pre-treatment) and after (post-treatment) hormonal manipulation in the different treatment groups (sham, metyrapone (mety) and cort). Pre-treatment are values for feathers plucked prior to hormonal manipulation (grew in the wild; fig. 1A) and used as reference values. In the post-treatment, "**New**" feathers grew during hormonal manipulation (feather P4, P7, S2, S4 and rectrix, right wing; Fig. 1B). "**Mature**" feathers were fully grown at the time of hormonal manipulation (feathers P2 and S5, left wing, Fig. 1C). Note that the reference values for the Mature feathers are from the right wing, while values of post-treatment are from the left wing.

| Feathers | **Pre-treatment** | | | **Post-treatment** | | |
|---|---|---|---|---|---|---|
| **New** | **mean** | **median** | **SD** | **mean** | **median** | **SD** |
| sham | 17.39 | 7.99 | 36.00 | 332.32 | 113.44 | 566.20 |
| mety | 24.64 | 7.5 | 102.60 | 378.56 | 205.7 | 480.93 |
| cort | 15.08 | 4.61 | 24.76 | 509.78 | 404.24 | 442.39 |
| **Mature** | | | | | | |
| sham | 10.65 | 4.69 | 14.91 | 1271.38 | 818.19 | 2040.85 |
| mety | 8.88 | 5.74 | 8.32 | 1653.27 | 1115.18 | 1046.44 |
| cort | 9.57 | 6.61 | 8.65 | 4906.05 | 4716.87 | 2387.39 |

### B. Further details of LC-MS/MS methods for feather CORT quantitation

SAMPLE EXTRACTION

Each feather sample had the calamus removed, and then was weighed and placed in a 13 × 100 mm culture test tube. Methanol (LC/MS grade, Fisher Scientific), internal standard solution (10 µM d8-corticosterone in methanol; CDN Isotope Inc) and calibrator (Steroids Inc) solutions (see Table A3) were cooled to 4˚C in an ice bath. 5 mL of cold methanol was pipetted into each test tube followed by adding 100 µL of the internal standard solution. The test tubes were then sealed with polyethylene caps and placed in a 4˚C fridge. After 20 hours, the feathers were removed from the test tubes. The extracts were dried down under N2 by use of Techne Sample Concentrator at 40˚C.

LC-APCI-MS/MS ANALYSIS

Dried feather extracts were reconstituted as 100 µL of $H_2O$/MeOH (50/50, v/v) for quantitation using an Agilent 1200 binary liquid chromatography (LC) system connected with an AB SCIEX QTRAP® 5500 tandem mass spectrometer (MS/MS) equipped with an atmospheric pressure chemical ionization (APCI) source. LC separation was performed on an Agilent Poroshell 120 C18 column (50 x 3 mm, 2.7 µm particle size) at 45˚C. The mobile phase A was $H_2O$/MeOH (75/25, v/v) and the mobile phase B was 100% methanol. The 8.2 min gradient was 20-70% B (0-3.8 min), 70-100% B (3.8-4.1 min), 100% B (4.1-5.5 min), 100-10% B (5.5-5.8 min), and held at 10% B (5.8-8.2 min). The flow rate was 0.6 mL/min and the injection volume was 10 µL.

Mass spectrometer conditions are listed in Table A4. Nitrogen was provided by an LC/MS 5500 nitrogen generator (Parker Balston) and was used as the source gas, nebulizer gas, and collision gas. Mass resolution in Q1 and Q3 was set to unit resolution. Two transitions were monitored for each analyte, a quantifier transition (-1) and a qualifier transition (-2), with conditions listed in Table A5.

### LOWER LIMIT OF QUANTITATION

A statistical analysis was used to estimate the lower limit of quantification (LLOQ) of corticosterone in feather samples.[1] The lowest concentration in which the CV is <20% and the error is <±20% was used as the LLOQ. All samples were prepared by spiking corticosterone standards into different matrices including pig saliva and stripped serum. Each sample was repeatedly injected ($n \geq 5$) and the accuracy and reproducibility were calculated. Details of the testing results are available as an xlsx file, on request.

For the feather extracts, three pools were created from sample injections that had been quantified as <0.1 ng/mL, 0.1–0.2 ng/mL and 0.2–0.3 ng/mL. Each was run in 10 replicates. The highest pool met the criteria for LLOQ at 0.25 ng/mL Samples between 0.1 and 0.2 ng/mL were statistically lower than the LLOQ and labeled as “< LOQ” in the results output. No samples fell below the detection limit. In a few samples, background was high, which decreased the reliability of the concentration. Samples with background ≥3500 counts were annotated as “high background”.

---

([1]) http://www.absciex.com/Documents/Downloads/Literature/mass-spectrometry-cms_059150.pdf

Table A3. Corticosterone concentrations (ng/mL) of calibration curve prepared in water. R2 of the calibration curve is > 0.99

| Compound | STD 1 | STD 2 | STD 3 | STD 4 | STD 5 | STD 6 | STD 7 | STD 8 | STD 9 |
|---|---|---|---|---|---|---|---|---|---|
| Corticosterone | 500 | 250 | 100 | 50 | 25 | 5 | 1 | 0.5 | 0.25 |

Table A4. Mass spectrometer conditions

| Parameter | Value |
|---|---|
| Curtain gas | 60 psi |
| Temperature | 500 °C |
| Ion Source Gas 1 | 50 psi |
| Collision Gas | Medium |
| Nebulizer Current | 5 µA |

Table A5. MRM conditions for corticosterone. (m/z= mass-to-charge ratio, DP= declustering potential, EP= entrance potential, CE= collision energy, and CXP= collision cell exit potential).

| Analyte | m/z of Parent Ion | m/z of Daughter Ion | DP (V) | EP (V) | CE (eV) | CXP (V) |
|---|---|---|---|---|---|---|
| Corticosterone-1 | 347.2 | 329.2 | 75 | 8 | 24 | 11 |
| Corticosterone-d8-1 | 355.3 | 337.3 | 75 | 8 | 24 | 11 |
| Corticosterone-2 | 347.2 | 121.1 | 75 | 8 | 32 | 11 |
| Corticosterone-d8-2 | 355.3 | 125.1 | 75 | 8 | 32 | 11 |

Table A6. Accuracies of QC samples.

| Sample name | Calculated Conc. (ng/mL) | Measured Conc. (ng/mL) | Accuracy (%) |
|---|---|---|---|
| Low QC | 4 | 3.68 | 91.897 |
| Low QC | 4 | 3.46 | 86.571 |
| Low QC | 4 | 4.15 | 103.76 |
| Low QC | 4 | 3.21 | 80.365 |
| Low QC | 4 | 3.41 | 85.174 |
| Med QC | 40 | 37.41 | 93.528 |
| Med QC | 40 | 33.53 | 83.818 |
| Med QC | 40 | 38.85 | 97.115 |
| Med QC | 40 | 40.20 | 100.51 |
| Med QC | 40 | 36.18 | 90.459 |
